# Integrated *in vivo* and transcriptomic analyses of lethal Oropouche virus infection reveal suppression of pathogenic host responses by antiviral therapy

**DOI:** 10.64898/2026.03.16.712021

**Authors:** Cássia Sousa Moraes, Gabriel Gonzalez, Akihiko Sato, Shigeru Miki, Atsuko Inoue, Koshiro Tabata, Joshua W. Kranrod, Chilekwa Frances Kabamba, Aiko Ohnuma, Keita Matsuno, Rio Harada, Shinji Saito, Michihito Sasaki, Yasuko Orba, William W. Hall, Hirofumi Sawa, Yukari Itakura

## Abstract

Oropouche virus (OROV) is an emerging arbovirus responsible for large outbreaks of febrile illness in Central and South America, with increasing reports of severe neurological disease and fatal outcomes. Despite its growing public health impact, no approved antiviral therapies or vaccines are currently available. Here, we show that favipiravir, a broad-spectrum nucleoside analogue, robustly suppresses OROV replication and disease *in vivo*. In a lethal Syrian hamster model, favipiravir treatment provided complete protection against OROV infection, preventing viral dissemination to peripheral organs and the central nervous system, and remained highly effective when administration was initiated after infection. In contrast, insufficient antiviral control resulted in viral neuroinvasion and fatality. To define host responses associated with OROV pathogenesis and their modulation by antiviral therapy, we performed transcriptomic profiling of liver and brain tissues. OROV infection induced interferon-driven inflammatory programs accompanied by marked disruption of metabolic and tissue homeostatic pathways, whereas these transcriptional signatures were largely abrogated by favipiravir treatment. Together, our findings identify favipiravir as a potent antiviral candidate against OROV and provide the first *in vivo*, tissue-resolved transcriptomic framework of OROV infection, linking effective viral suppression with the prevention of neuroinvasion and pathogenic host responses. These results highlight antiviral intervention as a viable strategy to mitigate OROV-associated disease and mortality.

## 1. Introduction

Oropouche virus (OROV) is an emerging arbovirus endemic to Central and South America which causes recurrent outbreaks of an acute febrile illnesses, collectively referred to as Oropouche fever^1^. Although most infections are self-limiting, an increasing number of cases have been associated with severe neurological manifestations, fatal outcomes, and adverse pregnancy events^2–8^. Since 2023, large-scale outbreaks involving tens of thousands of confirmed cases have been reported, including regions which had no prior evidence of OROV circulation, underscoring the escalating public health threat posed by this virus^1^. Despite the growing recognition of OROV as a clinically relevant pathogen, there are currently no approved antiviral therapies or vaccines, and clinical management remains entirely supportive.

OROV belongs to the genus *Orthobunyavirus* and can cause systemic infection with central nervous system (CNS) involvement^9^. Clinical reports and experimental studies have documented viral dissemination to peripheral organs and the brain, accompanied by neurological manifestations ranging from meningitis to encephalitis^7,8^. However, the mechanisms by which OROV infection leads to tissue damage, neuroinvasion, and severe disease remain poorly understood. In particular, it is unclear how viral replication interfaces with host antiviral, inflammatory, and metabolic pathways in different tissues to drive disease progression. Such interactions are critical determinants of disease outcome, as excessive or dysregulated host responses have been implicated in the pathogenesis of numerous neurotropic viral infections^10^.

A major limitation in this field is the lack of comprehensive *in vivo* analyses of host transcriptional responses during OROV infection. Although neuroinvasion and systemic disease have been demonstrated in both humans^8^ and animal models^11^, tissue-resolved transcriptomic studies which capture host responses in relevant target organs have not yet been reported. This gap has hindered a mechanistic understanding of OROV pathogenesis and limited the rational development of therapeutic interventions.

Favipiravir is a broad-spectrum nucleoside analogue that inhibits viral RNA-dependent RNA polymerases and is approved for the treatment of influenza in several countries^12,13^. This has demonstrated antiviral activity against a wide range of RNA viruses, including emerging and neurotropic pathogens^12,14–17^. Antiviral studies against OROV have so far been largely limited to *in vitro* discovery efforts. Broad-spectrum nucleoside analogues such as favipiravir, ribavirin and remdesivir inhibit viral replication in cultured cells, and recent screening approaches identified additional small-molecule and endonuclease-targeting inhibitors including acridones, lysergol, and 4′-fluorouridine^18–21^. Although *in vivo* antiviral activity has been reported^21^, the impact of favipiravir on systemic viral dissemination and host transcriptional responses during Oropouche virus infection has not been comprehensively investigated.

Here, we investigated the antiviral efficacy of favipiravir against OROV infection using *in vitro* assays and a lethal Syrian hamster model. We further performed tissue-resolved transcriptomic analyses of the liver and brain to define host responses associated with OROV pathogenesis and their modulation by antiviral therapy. We show that effective viral suppression by favipiravir prevents systemic dissemination and neuroinvasion, suppresses pathogenic inflammatory and metabolic transcriptional programs, and confers robust protection against lethal disease. Together, these findings establish favipiravir as a promising antiviral candidate against OROV and provide an *in vivo* framework that integrates viral replication with host responses and disease-associated pathways during infection.

## 2. Results

2.1. Favipiravir potently inhibits OROV replication *in vitro*

To evaluate the antiviral efficacy of favipiravir against OROV (TRVL 9760 strain), we performed an *in vitro* dose–response assays with Vero E6 cells using the MTT assays and compared its activity to ribavirin. Favipiravir showed a higher inhibitory effect, with mean half-maximal effective concentrations (EC_50_) of 28.4 µM, whereas ribavirin was less effective with EC_50_ of 115.7 µM (Fig. 1A). Consistently, ribavirin treatment produced only a modest decrease in virus titer across the tested concentrations. On the other hand, favipiravir exerted a robust, dose-dependent antiviral effect, reducing infectious OROV production (Fig. 1B). Untreated OROV-infected cells exhibited marked cytopathic effects (CPE), and ribavirin-treated cultures demonstrated only limited protection. In contrast, favipiravir treatment preserved cellular morphology, with a progressive reduction in CPEs as drug concentrations increased. (Fig. 1C).

**Fig 1.**
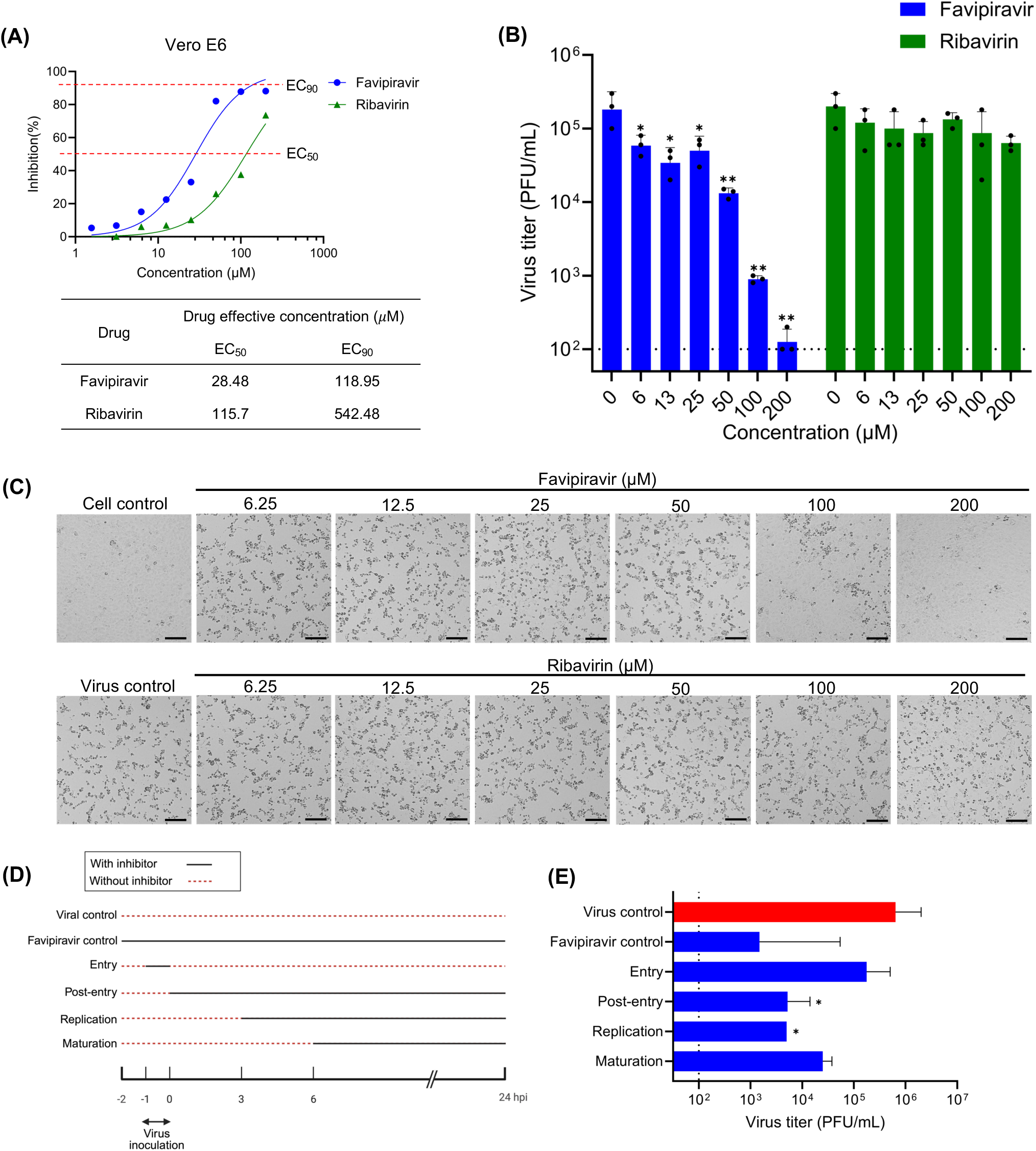
*In vitro* antiviral effects of favipiravir and ribavirin against OROV infection. (A-C) Vero E6 cells were infected with OROV TRVL 9760 at an MOI of 1. Favipiravir, or ribavirin was added 1 hour prior to infection. **(A)** Dose–response curves of antiviral activities of favipiravir and ribavirin were assessed by MTT assays at 72 hpi. Red dashed lines indicate 50% and 90% effective concentrations (ECLL and ECLL). **(B)** Virus titers in the supernatants were measured by plaque assay at 72 hpi. Statistical significance was assessed by two-way ANOVA with Tukey’s post-hoc test *(*p < 0.05, **p <0.01*). **(C)** Representative microscopy images of cells at 72 hpi. Scale bars: 100 µm. **(D, E)** Time-of-addition assays. Vero E6 cells were infected with OROV at an MOI of 1, and favipiravir (200 μM) was administered at the indicated time points. **(D)** Schematic image of the assay. **(E)** Virus titers in the supernatants at 24 hpi. Dashed lines indicate the limit of detection. Bars indicate the geometric means ± geometric standard deviation (s.d.) of three independent replicates. Statistical significance was assessed by one-way ANOVA with Dunnett’s post-hoc test *(*p < 0.05*).

To clarify which stage of the viral cycle is affected by favipiravir, we performed a time-of-addition assay (Fig. 1D). Cells exposed to the drug during the post-entry stage (0 hours post infection [hpi]), or the early replication stage (3 hpi) showed significantly reduced viral titer at 24 hpi compared with non-treated cells, whereas drug treatment during the maturation stage (6 hpi) did not exhibit a significant decrease in virus titers (Fig. 1E). These results demonstrated a strong inhibition after the onset of replication, consistent with favipiravir’s role as a nucleoside analogue that interferes with viral RNA synthesis, exerting its main antiviral effect during the post-entry replication phase.

Overall, these findings demonstrate that favipiravir efficiently inhibits OROV replication *in vitro* and offers superior protection from virus-induced CPEs compared to ribavirin.

### 2.2. Antiviral suppression by favipiravir prevents systemic dissemination and lethal OROV disease in Syrian hamsters

Given the broad antiviral activity of favipiravir as demonstrated in multiple experimental animal models, including viral hemorrhagic fever and encephalitic viral diseases^17^, we assessed its therapeutic efficacy against OROV infection employing a Syrian hamster infection model.

Animals were infected with a lethal dose of the OROV TRVL 9760 strain (5.4 × 10^5^ plaque-forming units [PFUs]) *via* the intraperitoneal route. Favipiravir (100 or 600 mg/kg/day) or ribavirin (40 or 100 mg/kg/day) was administered twice daily *via* the perioral route beginning at 1 hour before infection and continuing for 6 days (Fig. 2A).

**Fig 2.**
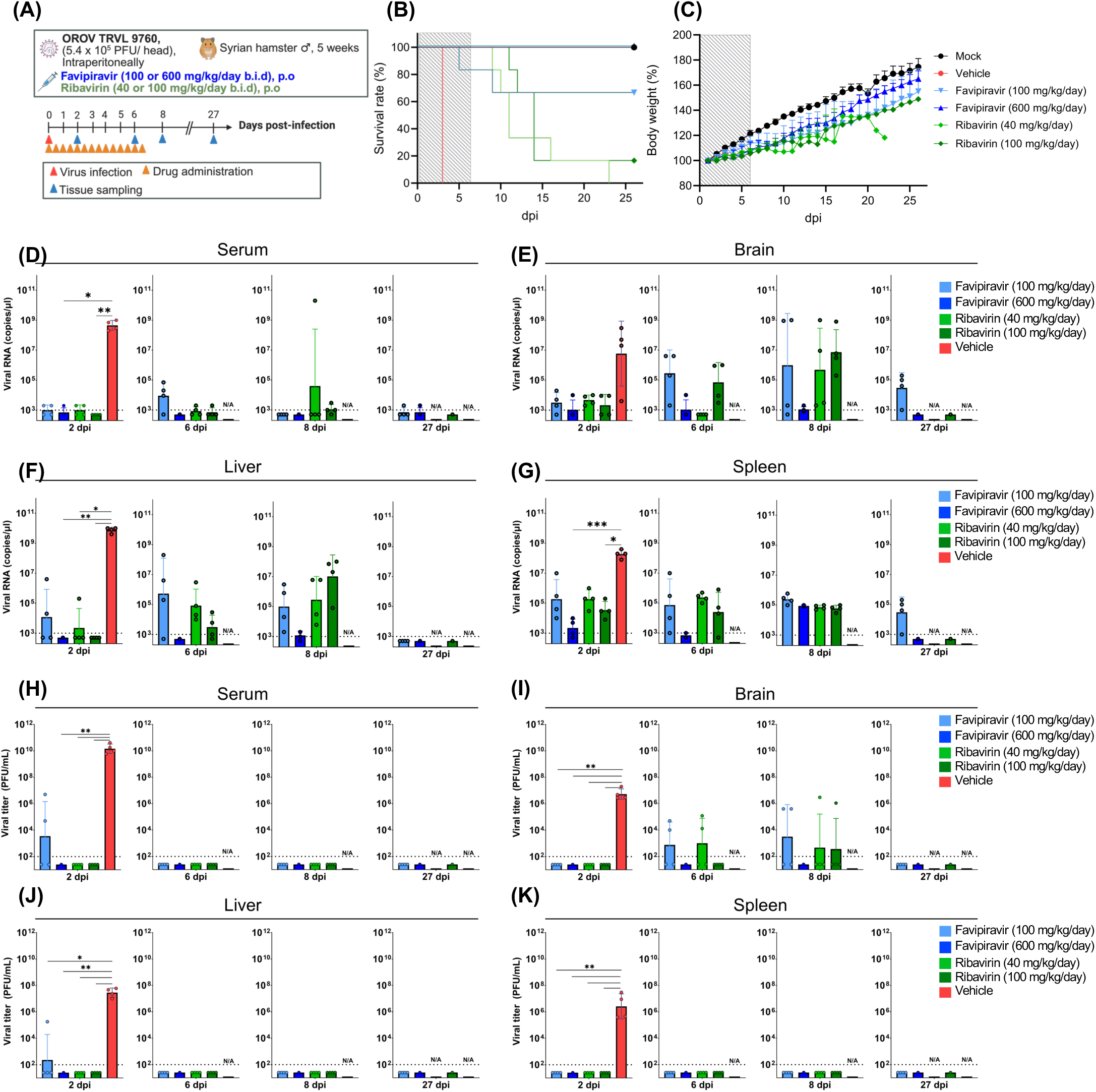
Prophylactic oral administration of favipiravir protects Syrian hamsters from lethal OROV infection. **(A)** Schematic illustration of the experimental design. Syrian hamsters were infected intraperitoneally with OROV (5.4 x 10^5^ PFU/ head) and treated with favipiravir (100 or 600 mg/kg/day) or ribavirin (40 or 100 mg/kg/day) starting 1 hour before infection, twice daily until 6 days post-infection (dpi). The dashed grey box indicates the treatment window. Animals were monitored for **(B)** survival and **(C)** body weight changes until 26 dpi. **(D-G)** Quantification of viral RNA by RT-qPCR in the **(D)** serum, **(E)** brain, **(F)** liver and **(G)** spleen collected at the indicated time points. Copy numbers were calculated based on standard curves. **(H**-**K)** Infectious viral titers determined by plaque assay in the **(H)** serum, **(I)** brain, **(J)** liver and **(K)** spleen at the indicated time points. **(D-K)** Vehicle-treated animals succumbed to infection after 2 dpi; therefore, samples were only available at 2 dpi for this group. N/A: not applicable. Dashed lines indicate the limit of detection. Bars represent geometric means ± geometric s.d. and dots represent individual animals. Statistical significance was performed by Kruskal-Wallis followed by Dunn’s multiple comparison post-hoc test: *\*p< 0.05, **p<0.01 and ***p<0.001*.

**Fig 3.**
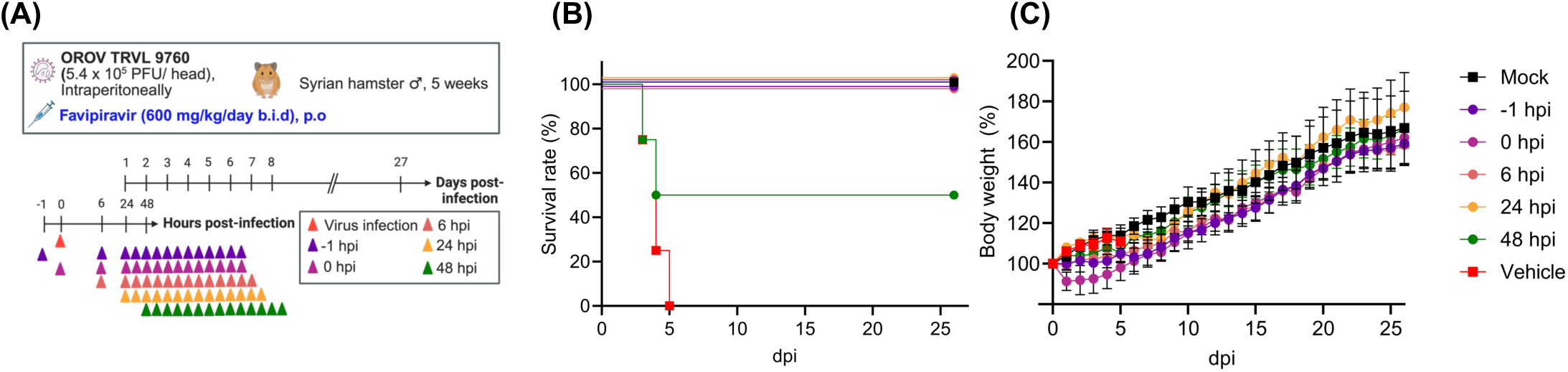
Delayed favipiravir treatment protects Syrian hamsters from lethal OROV infection. (**A)** Experimental design of the delayed-treatment model. Syrian hamsters were infected intraperitoneally with 5.4L×L10LLPFU of OROV and received favipiravir (600Lmg/kg/day, orally, twice daily for 6Ldays) starting at the indicated time points. **(B)** Survival and **(C)** body weight was monitored for 26Ldays.

All vehicle-treated animals succumbed to infection by 3 days post-infection (dpi) without preceding overt clinical symptoms or body weight loss. In contrast, animals treated with high-dose favipiravir (600 mg/kg/day) survived the 26-day study period without exhibiting any clinical signs and steadily gained weight throughout the trial. Treatment with a low dose of favipiravir (100 mg/kg/day) or ribavirin (40 or 100 mg/kg/day) conferred partial protection (Fig. 2B and 2C). Notably, following treatment cessation at 6 dpi, some animals in each of the ribavirin groups exhibited weight loss and developed neurological symptoms, including lethargy, shivering, and hind-limb paralysis (S1 Fig.).

Viral replication in tissues and sera at 2, 6, 8, and 27 dpi was assessed by plaque assay and RT-qPCR. While high viral loads were detected in all tissues and sera of vehicle-treated animals at 2 dpi, no viral RNA or infectious virus was detected in animals treated with high-dose favipiravir at any time point across all tissues (Fig. 2D-K). Treatment with low-dose favipiravir or ribavirin effectively reduced infectious virus production in serum, liver, and spleen (Fig. 2H, 2J, and 2K). Importantly, infectious virus was detected in the brains of these groups at 6 and 8 dpi (Fig. 2I), indicating incomplete clearance of OROV from the CNS at suboptimal dosing, and which was associated with the neurological symptoms observed after treatment cessation.

Collectively, these results demonstrate that favipiravir provides complete protection against lethal OROV infection in hamsters, suppressing systemic viral replication and limiting viral neuroinvasion.

### 2.3. Therapeutic administration of favipiravir after infection blocks viral neuroinvasion and improves surviva**l**

To assess the therapeutic potential of favipiravir, we examined its efficacy when treatment was delayed following OROV infection. Hamsters infected with OROV were administered favipiravir (600 mg/kg/day) starting at –1, 0, 6, 24, or 48Lhpi (Fig.L3A). All animals receiving favipiravir from –1, 0, 6, and 24Lhpi were fully protected, with 100% survival (Fig.L3B). Remarkably, even when administration was initiated as late as 48Lhpi, favipiravir still conferred partial protection, with 50% survival and consistent weight gain during the observation period (Fig.L3B-C and S2 Fig.).

These findings highlight favipiravir’s potential for clinical use when treatment cannot always be initiated immediately post-exposure.

### 2.4. Tissue-resolved transcriptomic analyses reveal pathogenic host programs induced by OROV infection and abrogated by favipiravir

The mechanisms underlying OROV-induced disease remain poorly characterized in both human cases and animal models. To define OROV infection–associated host responses and determine how favipiravir modulates these processes, we performed RNA sequencing (RNA-seq) of liver and brain tissues collected at 2 dpi, guided by the *in vivo* observation of hepatic and neurological involvement. Principal component analysis (PCA) demonstrated a clear separation between uninfected and vehicle-treated infected animals in both tissues (Fig. 4A-B). In the liver, one infected animal with low viral replication clustered closer to uninfected animals (Fig. 4A-B and S3 Fig.). Importantly, favipiravir-treated animals clustered near the uninfected control group, indicating that marked infection-driven transcriptional changes were largely suppressed by favipiravir treatment.

**Fig 4.**
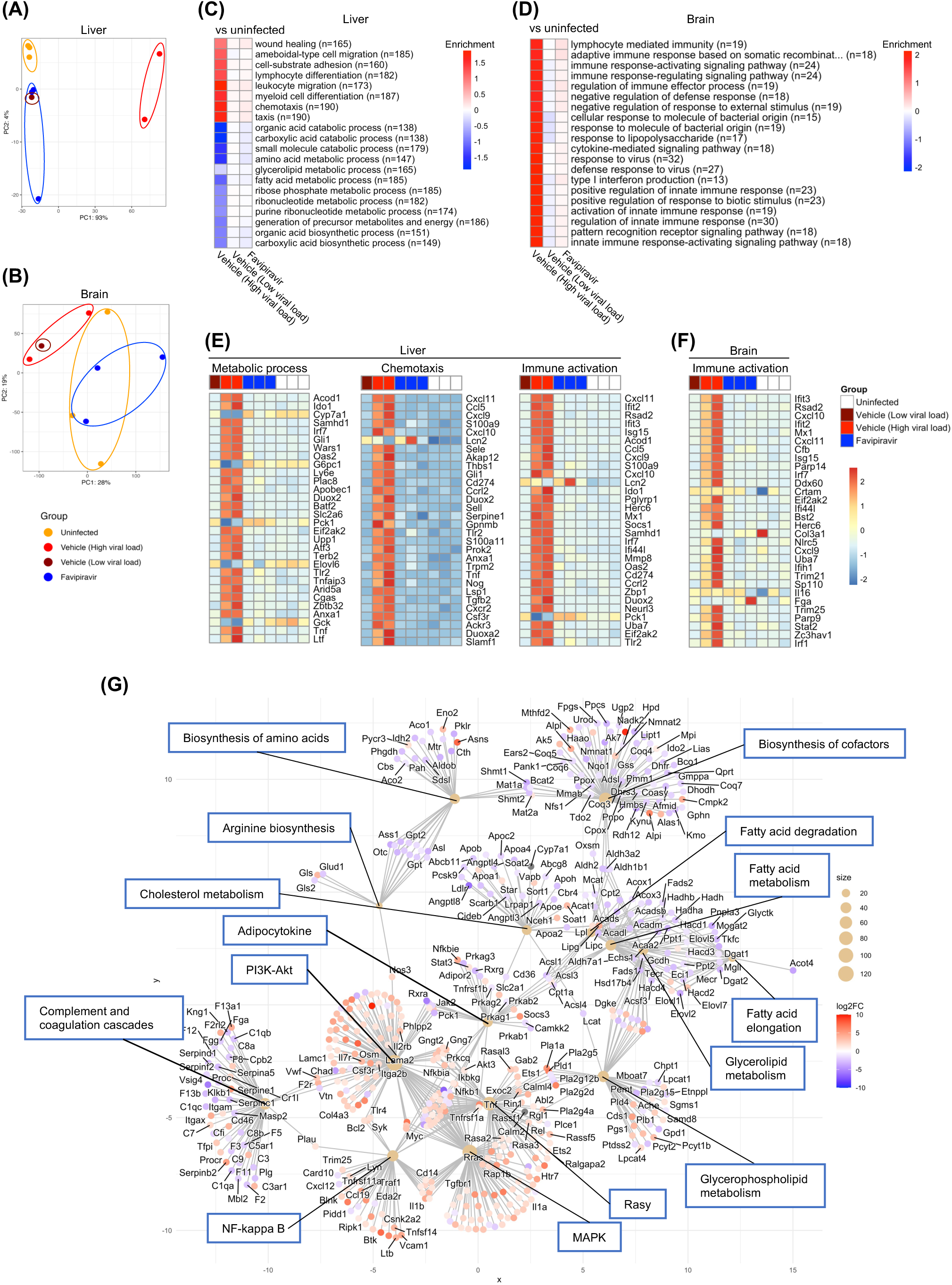
OROV infection alters host transcriptional responses in the liver and brain and favipiravir reversed these changes. (A-F) The liver and brain at 2 dpi were subjected to the RNAseq analysis. Principal component analysis (PCA) in the liver **(A)** and brain **(B)**. Gene ontology (GO) enrichment analysis of differentially expressed genes (DEGs) in the liver **(C)** and brain **(D)**. The n represents the number of genes related to each pathway that are differentially expressed. Heatmaps of standardized expression levels of genes in the selected metabolic processes in the liver **(E)** and brain **(F)**. KEGG pathway–gene network analysis of the hepatic transcriptional response to OROV infection **(G)**.

Gene ontology (GO) enrichment analyses revealed that OROV infection induced robust systemic inflammatory and antiviral responses. In the liver, high viral replication induced upregulation of biological processes related to chemotaxis, cytokine signaling, myeloid cell differentiation, and interferon responses, along with downregulation of metabolic pathways related to energy production and fatty acid metabolism (Fig. 4C, 4E, 4G, and S4A Fig.). In the brain, antiviral and inflammatory pathways were more strongly induced, and accompanied by the promotion of immune cell recruitment (Fig. 4D, 4F, and S4B Fig.). These transcriptional modulations were markedly attenuated in favipiravir-treated animals, which maintained expression profiles similar to baseline in both tissues (Fig. 4C-F, and S4 Fig.).

To validate the RNA-seq findings and assess time-course changes, we quantified the expression of selected inflammatory and IFN-related genes (*Cxcl10, Rsad2,* and *Isg15*) in the liver (Fig. 5A) and brain (Fig. 5B) at 2, 6, and 8 dpi by RT-qPCR. At 2 dpi, expression of these genes was strongly induced in vehicle-treated animals in both tissues, whereas both favipiravir (600 mg/kg/day) and ribavirin (100 mg/kg/day) treatment suppressed this early induction. At 6 and 8 dpi, *Cxcl10* expression in both antiviral-treated groups remained at levels comparable to that in mock-infected animals. In contrast, expression of *Rsad2* and *Isg15* in brain and liver tissues tended to remain elevated at 6 and 8 dpi in ribavirin-treated animals, correlating with increased viral loads (Fig. 2E and 2I), whereas it remained suppressed in favipiravir-treated animals.

**Fig 5.**
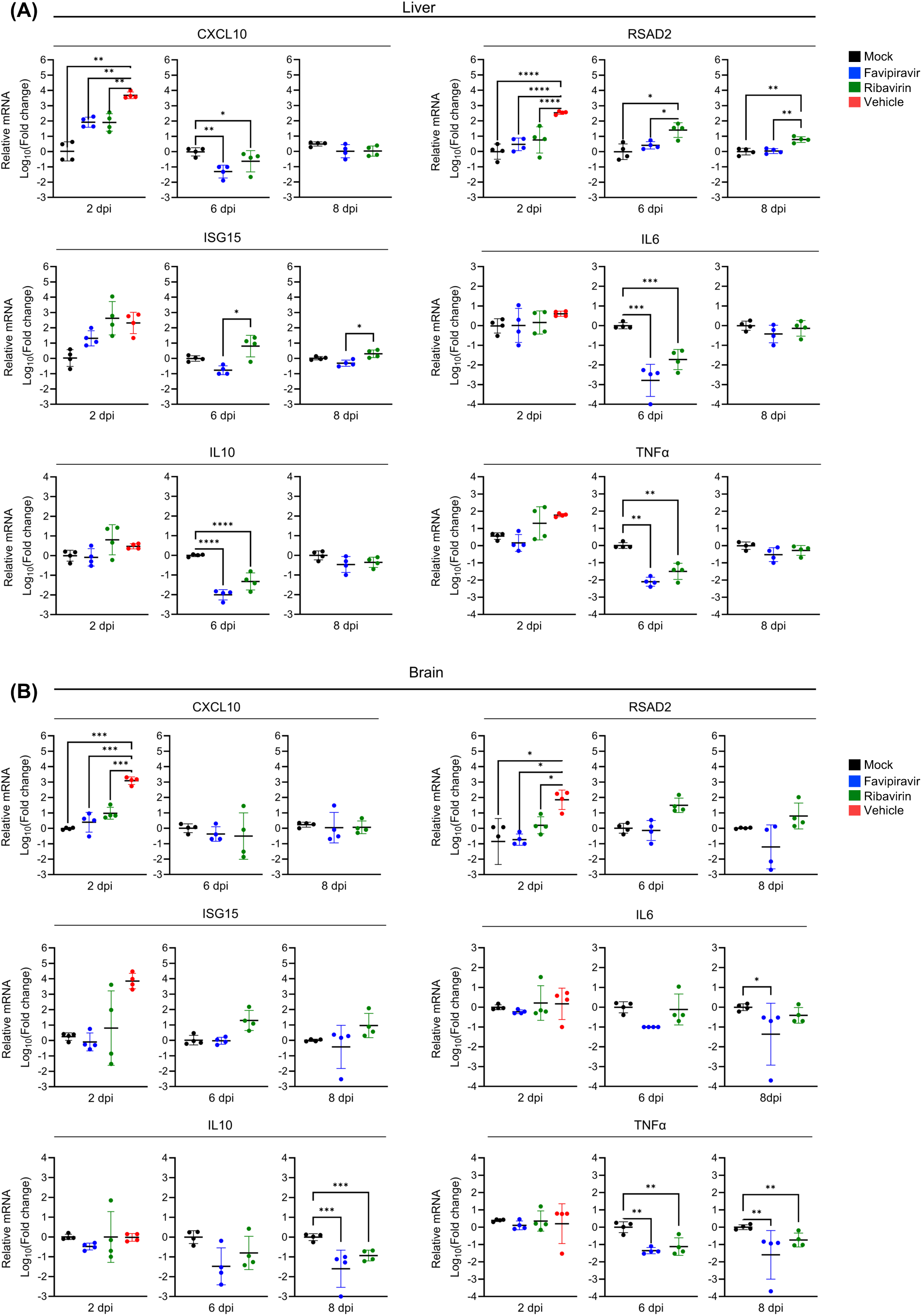
Validation of differential gene expression associated with immune and inflammatory responses. Relative gene expression of cytokines and interferon-stimulated genes in the liver **(A)** and **(B)** brain of hamsters at 2, 6 and 8 dpi. mRNA expression was quantified using the ΔΔCt method. Data were normalized to the housekeeping gene *Actb* and presented as fold changes relative to mock-animals. All graphs show geometric means ± geometric s.d. of 4 independent animals. Statistics were calculated by one-way ANOVA with Tukey’s post-hoc test: *\*p< 0.05, **p<0.01, and ***p<0.001 and ****p<0.0001*.

To further characterize host inflammatory responses during OROV infection, we evaluated the expression of *Il6, Il10* and *Tnf*α, three canonical inflammatory cytokines. At 2 dpi, none of these cytokines showed significant changes in either liver or brain tissues across the experimental groups. Although their expression decreased at later stages in both favipiravir-and ribavirin-treated animals, these changes did not correlate with viral replication, suggesting that these cytokines are unlikely to be major drivers of OROV-associated disease in this model.

Collectively, these data indicate that favipiravir provides sustained suppression of OROV-induced inflammatory and IFN responses in both the liver and the brain, whereas incomplete viral control by ribavirin is associated with re-emergence of IFN-stimulated gene expression at later stages of infection.

## 3. Discussion

OROV has historically been regarded as a cause of a range of self-limited febrile illnesses; however, recent outbreaks, confirmed human fatalities, and increasing reports of neurological and obstetric complications have redefined OROV as an emerging pathogen of significant public health concern^22–25^. Despite this growing clinical relevance, the mechanisms underlying severe disease and associated host responses remain poorly characterized, and no approved antiviral treatments are currently available. In this study, we demonstrate that favipiravir exhibits potent antiviral activity against OROV *in vitro* and provides robust therapeutic protection in a lethal Syrian hamster model. Importantly, our integrated virological and transcriptomic analyses have defined host response pathways associated with OROV infection and show that these disease-linked responses are effectively attenuated by antiviral treatment.

Favipiravir showed greater antiviral efficacy than ribavirin *in vitro*, with a substantially lower EC_50_ and strong suppression of OROV-induced CPE (Fig. 1A-C). Time-of-addition experiments indicated that favipiravir acts predominantly during the post-entry replication phase (Fig. 1D-E), consistent with its established mechanism as a nucleoside analogue targeting the viral RNA-dependent RNA polymerase^14,16^.

*In vivo*, favipiravir (600 mg/kg/day) conferred complete protection against lethal OROV challenge in hamsters, even when administration was initiated at 24 hpi, fully suppressing viral replication in systemic organs and the CNS (Fig. 2A-K and 3A-C). In contrast, suboptimal antiviral treatment or ribavirin administration resulted in incomplete viral clearance from the brain (Fig. 2I) and was associated with delayed onset of neurological signs alongside relative increases in inflammatory responses (Fig. 5B). Disease progression, including motor impairment and, in severe cases, paralysis, is consistent with the hematogenous spread of the virus to neural tissues, as suggested by previous animal studies^11,25^. These findings parallel human clinical observations in which neurological manifestations occur in approximately 4% of clinical cases^26^ along with reports of OROV RNA detection in the cerebrospinal fluids^8,27^. Collectively, these data suggest that timely and potent antiviral intervention by favipiravir may prevent CNS invasion and subsequent neurological disease during OROV infection, highlighting a critical therapeutic window relevant to human disease.

Although liver involvement is not a prominent feature of typical Oropouche fever, emerging clinical evidence indicates that hepatic dysfunction may occur in severe cases. Recent fatal OROV infections have been associated with markedly elevated aminotransferase levels, liver injury biomarkers^7^, suggesting that hepatic infection may contribute to disease pathogenesis in humans. In the present study, OROV exhibited robust viral replication in the liver, accompanied by strong inflammatory and interferon-driven responses, as well as suppression of metabolic and mitochondrial pathways (Fig. 4C, 4E, 4G, and 5A). These alterations were effectively normalized by favipiravir treatment, consistent with complete suppression of viral replication (Fig. 2F, 2J, and 4C). Although species-specific differences may partly explain the limited recognition of liver involvement in human OROV infection, hepatic infection may be associated with systemic inflammation and metabolic dysregulation in severe disease settings^28–30^, as suggested by this animal model, and favipiravir could represent a therapeutic option for such conditions with hepatic involvement.

To our knowledge, this study provides one of the first comprehensive *in vivo* characterizations of host transcriptional responses to OROV infection and identifies pathways likely linked to systemic inflammation related to disease severity. Transcriptomic analyses revealed robust innate immune activation in both liver and brain, including IFN-stimulated gene expression, inflammatory signaling, and immune cell recruitment, accompanied by acute disruption of energetic and lipid metabolic pathways in the liver and neuroinflammation in the brain (Fig. 4C-F and S4 Fig.). Notably, favipiravir limited virus-driven inflammation and preserved metabolic and homeostatic pathways (Fig. 4C-F and 5A-B), indicating that effective viral control prevents the activation of downstream pathogenic host responses. Together, these findings provide a mechanistic context for severe OROV disease, thereby advancing understanding of its pathogenesis, and suggest that the therapeutic benefit of favipiravir extends beyond viral load reduction to include modulation of host pathways associated with disease severity.

Recent studies have demonstrated the efficacy of another polymerase-targeting antiviral, 4′-fluorouridine against OROV *in vivo*^21,31^. Favipiravir offers distinct translational advantages, including oral bioavailability, an established safety profile, and prior clinical use against multiple viral diseases^14,32^. Importantly, the observed efficacy of delayed treatment initiation enhances its relevance for outbreak settings, where early diagnosis and immediate therapy may not always be feasible (Fig. 3B).

In conclusion, this study has established favipiravir as a potent antiviral candidate against OROV and provides one of the first *in vivo* frameworks linking viral replication, host transcriptional responses, and disease-associated pathways in OROV infection. Our findings provide strong preclinical evidence that favipiravir can effectively suppress systemic and neuroinvasive viral replication while preventing the activation of inflammatory and metabolic pathways associated with pathology. Our findings advance understanding of OROV pathogenesis and support the potential repurposing of favipiravir as a targeted therapeutic option for Oropouche fever.

## 4. Materials and Methods

### 4.1. Study design

This study was designed to evaluate the antiviral drugs against OROV infections. All viral experiments were performed in a biosafety level 3 laboratory at the International Institute for Zoonosis Control, Hokkaido University. Animal experiments with viral infection were performed according to the National University Corporation, Hokkaido University Regulations on Animal Experimentation. The protocol was reviewed and approved by the Institutional Animal Care and Use Committee of Hokkaido University (approval no. 24-0034). Experiments were done at least twice. All experiments with biological replicates are indicated in the figure legends.

### 4.2. Cells and viruses

Vero E6 (African green monkey kidney, clone E6) and BHK-21 cells were cultured and maintained in Dulbecco’s Modified Eagle’s Medium (DMEM; Gibco) supplemented with 10% heat-inactivated fetal bovine serum (FBS) at 37°C in a humidified incubator with 5% CO_2_. OROV TRVL 9760 strain was kindly provided by World Reference Center for Emerging Viruses and Arboviruses (WRCEVA), University of Texas Medical Branch at Galveston, UTMB Health and propagated in Vero E6 cells, and working virus stocks were stored at −80°C until use.

### 4.3. Compounds

For *in vitro* studies, favipiravir (FUJIFILM Wako Pure Chemical Corporation) and ribavirin (Tokyo Chemical Industry Co. Ltd.) were dissolved in dimethyl sulfoxide (DMSO; Sigma-Aldrich). For *in vivo* administration, favipiravir was prepared in 0.5% (w/v) carboxymethylcellulose (CMC; Sigma-Aldrich) and delivered via oral gavage. Ribavirin was dissolved in a vehicle solution containing 5% DMSO in 0.5% CMC.

### 4.4. MTT assays and EC_50_ determination

Cell viability was assessed using the 3-(4.5-dimethylthiazol-2yl)-2,5diphenyltetrazolium bromide (MTT) assay. Cells and virus were diluted in MEM supplemented with 2% FBS and added simultaneously with compounds in 96-well plates. Briefly, 50 µL of two-fold serially diluted favipiravir or ribavirin was dispensed into each well, followed by 100 µL of cell suspension (2.5 × 10L cells) and 50 µL of virus suspension. The amount of virus used corresponded to approximately 4 TCIDLL. Plates were incubated at 37 °C for 72 h. After incubation, 30 µl of MTT solution (5 mg/L) was added, and the plates were incubated at 37°C for 4 hours. Then, the supernatant was removed, and 140 µl of MTT solubilizer buffer was added to dissolve the formazan crystal overnight at room temperature. MTT reactions were measured using a microplate reader at an optical density (OD) of 570 nm. The results are presented as the percentage of viable cells relative to untreated control cells. The 50% effective concentration (EC_50_) was determined using logarithmic interpolation (% of inhibition was calculated as (OD_sample_L−LOD_virus control_)/(OD_cell control_L−LOD_virus control_)) using GraphPad Prism with a four-parameter curve-fitting analysis.

### 4.5. Time of addition assay

Vero E6 cells were infected with OROV at a MOI of 1. Following virus adsorption for 1 hour at 37°C, cells were washed with PBS and 2% FBS DMEM was added. Favipiravir (50 µM) was added at: 1 hour before infection (−1 hpi), at the time of infection (0 hpi), and at 1-, 3-, and 6-hpi. Supernatants were harvested at 24 hpi, and viral replication was quantified by plaque assay.

### 4.6. Plaque assays

Monolayers of Vero E6 were inoculated with serial ten-fold dilutions of the specimens and incubated for 1 hour at 37°C. After incubation, cells were overlaid with 0.5% methyl cellulose in Eagle’s Minimum Essential Medium (EMEM, Nissui Pharmaceutical Co., Ltd.) supplemented with 5% FBS and GlutaMAX supplement (Gibco). After 3 days of incubation, cells were fixed with a neutral 10% formalin neutral buffer solution and stained with 1% crystal violet to visualize plaques. Virus titers were calculated as PFU per milliliter (PFU/mL).

### 4.7. *In vivo* antiviral efficacy assessment in hamsters

Five-week-old male Syrian hamsters were intraperitoneally infected with 5.4 × 10^5^ PFU of the OROV TRVL 9760 strain in 200 µL of PBS. Favipiravir (100 or 600 mg/kg/day) or ribavirin (40 or 100 mg/kg/day) was administered orally beginning 1 hour before infection (-1 hpi) and continued twice daily until 6 dpi. The doses of favipiravir and ribavirin were selected based on previous pharmacokinetic and safety data in hamsters, in which no toxicity was observed^40–44^. For survival experiments, animals were monitored daily for symptoms and bodyweight changes up to 27 dpi. For tissue or serum sample collection, animals were euthanized at 2, 6, 8 dpi, with survivors sacrificed at 27 dpi, and the brain, liver, spleen, and serum were collected. Tissues were homogenized in either in PBS, for viral titration and RT-qPCR or RNA Shield (Zymo Research) for RNAseq. For delayed treatment with favipiravir, hamsters were orally administered 600 mg/kg/day of favipiravir twice daily, starting at −1, 0, 6, 24, or 48 hpi. The humane endpoints were defined by daily observation of severe clinical symptoms or weight loss exceeding 30%.

### 4.8. Reverse Transcription - quantitative PCR (RT-qPCR) assay

Total RNA was extracted from hamster tissue homogenates using TRIzol LS reagent (Invitrogen) and Direct-zol RNA Miniprep kit (Zymo Research) according to the manufacturer’s instructions. RT-qPCR targeting OROV was performed using Thunderbird Probe One-step RT-qPCR kit (Toyobo) following the manufacturer’s recommended cycling conditions: 50°C for 10 minutes, 95°C for 1 minute, followed by 45 cycles of 15 seconds at 95°C and 45 seconds at 60°C. Reactions were carried out on a LightCycler 96 Real Time System (Roche). Primers and probe targeting the S segment of OROV genome were designed as previously described (S1 Table). The viral copy number was estimated by the standard curve method.

Measurements of host gene expression was performed using One Step TB Green PrimeScript RT-PCR Kit II (Perfect Real Time) (Takara). Primers for hamster *Actb, Cxcl10, Il6, Il10, Isg15, RSAD2 and Tnf*Cl, are listed in the S1 Table. Target mRNA levels were normalized to hamster *Actb* mRNA and calculated as fold change relative to mock infected samples with the delta-delta cycle threshold (ΔΔCT) method.

### 4.9. Transcriptomic analysis

For mRNA-seq analysis, livers and brains at 2 dpi were stored in RNA Shield for 24 hours, then total RNA was extracted and purified using TRIzol LS reagent (Invitrogen), chloroform, and Direct-Zol RNA Miniprep kit (Zymo Research) according to the manufacture’s protocol. Sequencing libraries were prepared using NEBNext Ultra II Directional RNA Library Prep Kit for Illumina (New England BioLabs) and sequenced on an Illumina NovaSeq with 150 base pair-end reads at Azenta Life Science (S2 Table). Reads were processed with Trim Galore v0.6.10 for quality control, trimming and filtering. Preprocessed paired reads were aligned to a reference genome (accession number: GCA_017639785.1, *Mesocricetus auratus*) with Bowtie v2. RNA-seq counts data were filtered out to remove genes with low counts (<10 reads). RNA-seq data were analyzed in R using DESeq2 version 1.44.0 for normalization and differential expression. PCA was applied to normalized gene expression data to identify groups of samples that behave similarly or exhibit similar characteristics using the R package DeSeq2, and the results were visualized with *ggplot2* 4.0.0. Functional enrichment analysis was conducted with *clusterProfiler* version 4.12.6^33^. Due to incomplete annotation of *Mesocricetus auratus*, differentially expressed hamster gene symbols were mapped to their *Mus musculus* ortholog gene symbols prior enrichment analysis. The conversion was done via Biomart (Ensembl) mapping. Any gene lacking a valid mouse ortholog was removed from the downstream analysis. Over-representation analysis (ORA) was performed using GO. For ORA, significance was determined with *P* values <0.05 (adjusted by the Benjamini-Hochberg method) of a one-sided version of Fisher’s exact test. Heatmaps were generated with the R package *pheatmap* version 1.0.13. Kyoto Encyclopedia of Genes and Genomes (KEGG) pathway enrichment analysis was performed using the enrichKEGG function from the clusterProfiler R package, using the mapped gene set described above. Genes with adjusted *p* values < 0.05 and an absolute log_2_ fold change > 1 were included in the enrichment analysis. KEGG pathways with adjusted *p* values < 0.05 were considered significantly enriched. Network-based visualization of enriched KEGG pathways and their associated genes was generated using the cnetplot function from the enrichplot package version 1.30.4.

### 4.10. Statistical analysis

All statistical analyses were performed using Prism version 10.4.2 (GraphPad Software). Comparisons of RNA copies RT-qPCR and quantification of viable virus were performed using *Kruskal-Wallis* test followed by Dunn’s multiple comparison post-hoc test. *P* values of ≤ 0.05 were considered significant. For the validation of host expression genes, statistical significance of the fold difference between the infected and the uninfected samples was calculated with the ordinary one-way ANOVA followed by Tukey’s post-hoc test.

Transcriptomic data were analyzed and plotted in R version 4.4.0, using the above-mentioned R packages.

### 4.11. Data availability

Raw sequencing data were deposited at the NCBI Sequence Read Archive (SRA) under BioProject accession number PRJNA1416565.

## Acknowledgments

We would like to thank Mr. Kei Konishi at the International Institute for Zoonosis Control, Hokkaido University, for the helpful advice on the preparation of the antiviral drugs used in the *in vivo* analysis. We sincerely appreciate WRCEVA, UTMB Health for providing OROV. This study was supported in part by the Japan Society for the Promotion of Science (JSPS) KAKENHI under grant numbers 25K18810 and 25K09431; and the Japan Agency for Medical Research and Development (AMED) under grant numbers JP223fa627005 and JP24wm0225044. Schematic figures were created with BioRender.com.

## Author Contributions

**Conceptualization:** Cássia Sousa Moraes, Yukari Itakura

**Data Curation:** Cássia Sousa Moraes, Gabriel Gonzalez, Yukari Itakura

**Formal Analysis:** Cássia Sousa Moraes, Gabriel Gonzalez, Yukari Itakura

**Funding Acquisition:** Hirofumi Sawa, Yukari Itakura

**Investigation:** Cássia Sousa Moraes, Atsuko Inoue, Koshiro Tabata, Akihito Sato, Shigeru Miki, Joshua W. Kranrod, Chilekwa Frances Kabamba, Rio Harada, Yukari Itakura

**Methodology:** Cássia Sousa Moraes, Gabriel Gonzalez, Yukari Itakura

**Project Administration:** Hirofumi Sawa, Yukari Itakura

**Resources:** Akihito Sato, Keita Matsuno, Michihito Sasaki, Yasuko Orba, Aiko Ohnuma, Hirofumi Sawa, Yukari Itakura.

**Supervision:** Hirofumi Sawa, Yukari Itakura

**Validation:** Cássia Sousa Moraes

**Visualization:** Cássia Sousa Moraes

**Writing – Original Draft Preparation:** Cássia Sousa Moraes

**Writing – Review & Editing:** Cássia Sousa Moraes, Gabriel Gonzalez, Akihiko Sato, Shigeru Miki, Atsuko Inoue, Koshiro Tabata, Joshua W. Kranrod, Chilekwa Frances Kabamba, Aiko Ohnuma, Keita Matsuno, Rio Harada, Shinji Saito, Michihito Sasaki, Yasuko Orba, William W. Hall, Hirofumi Sawa, Yukari Itakura.

