## Supplementary Figures for "Integrated *in vivo* and transcriptomic analyses of lethal Oropouche virus infection reveal suppression of pathogenic host responses by antiviral therapy"

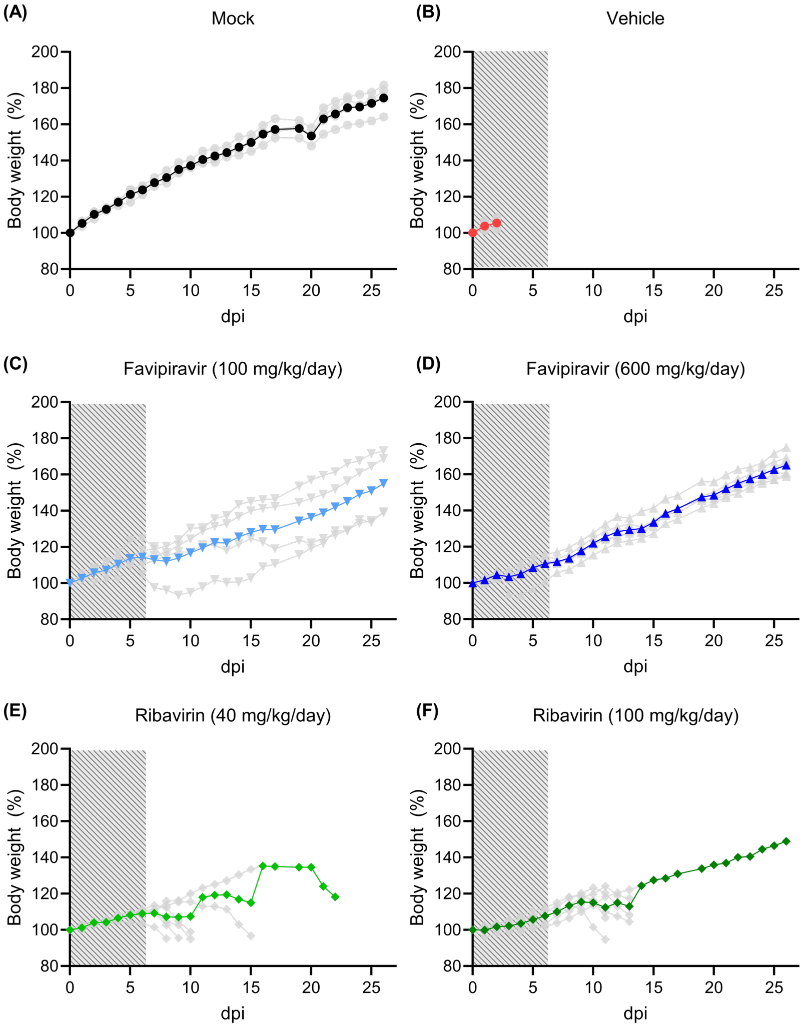


**S1 Fig. Individual body weight changes of hamsters in Fig. 2C**. Bold lines represent the average of the body weight in each group while grey lines represent individual body weight values in **(A)** mock, **(B)** vehicle, **(C-D)** favipiravir (100 or 600 mg/kg/day), and **(E-F)** ribavirin (40 or 100 mg/kg/day) groups. Dashed grey box indicates the treatment window **(B-F)**.

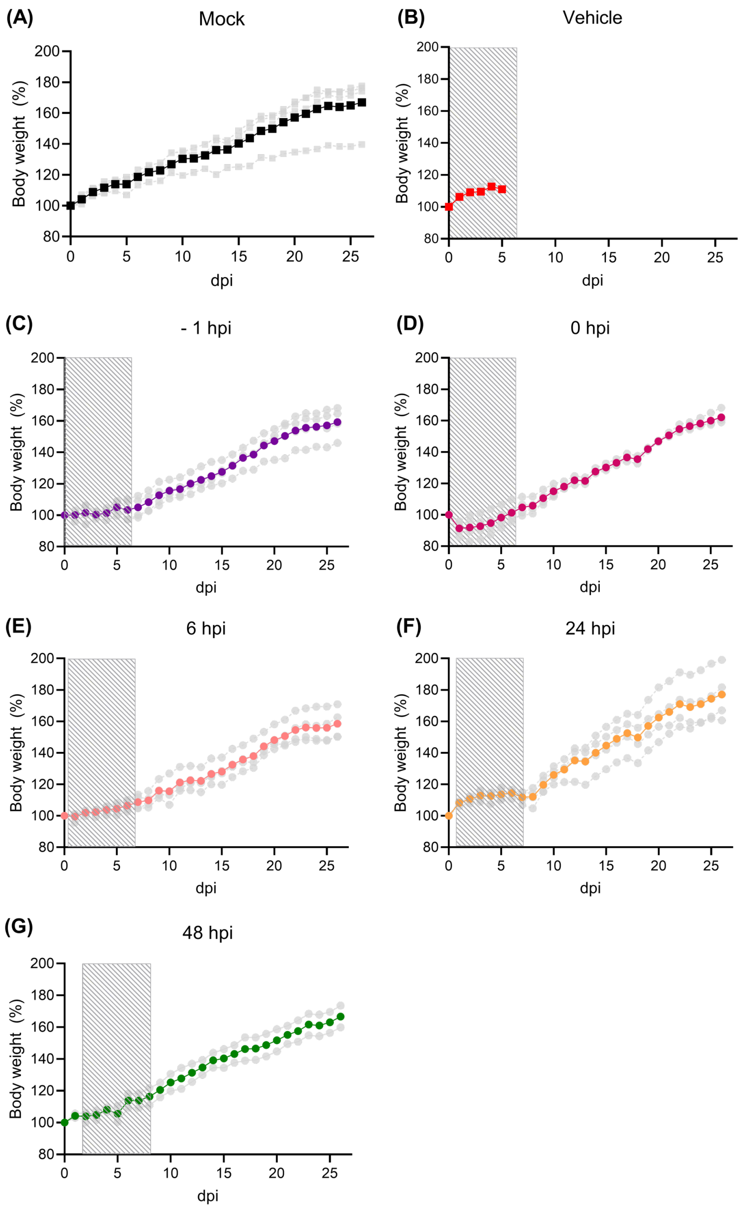


**S2 Fig. Individual body weight changes of hamsters in Fig. 3C**. Bold lines represent the average of the body weight in each group while grey lines represent individual body weight values in **(A)** mock, **(B)** vehicle, **(C)** -1 hour before infection (-1 hpi), **(D)** at the time of infection (0 hpi), **(E)** 6 hpi, **(F)** 24 hpi, and **(G)** 48 hpi groups. Dashed grey box indicates the treatment window **(B-G)**.

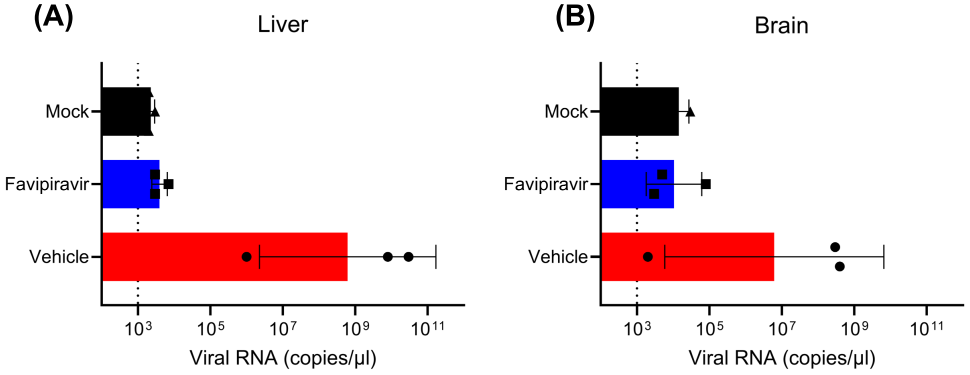


**S3 Fig. Viral RNA copies in the liver and brain used for RNAseq analysis.** Viral RNA copies were measured based on the standard curve by RT-qPCR in the brain **(A)** and liver **(B)** of hamsters at 2 dpi. Dashed lines represent the limit of detection. Bars represent geometric mean ± geometric s.d ., with individual biological replicates shown as dots

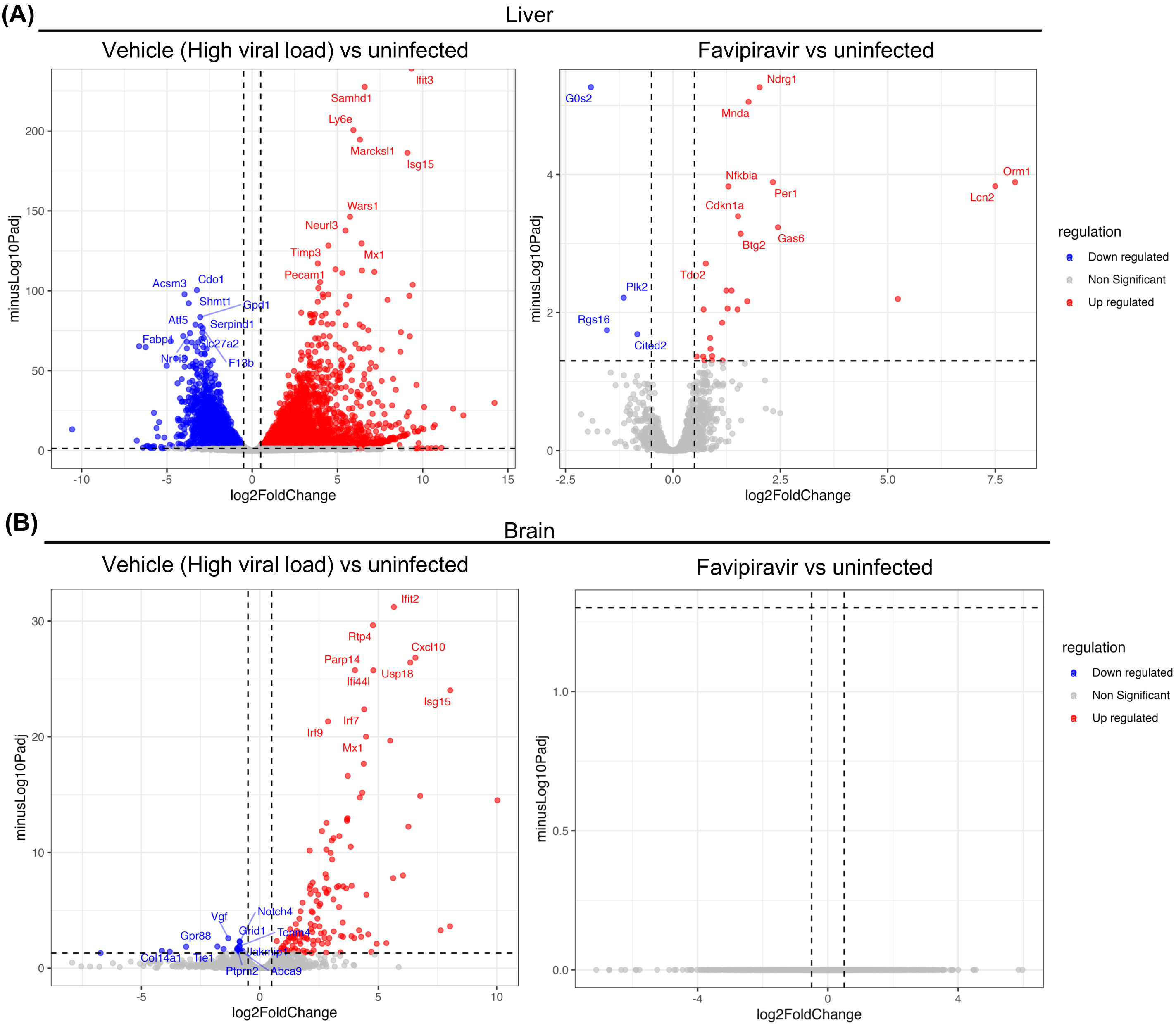


**S4 Fig. Transcriptomic profiling of OROV-infected hamsters related to Fig. 4.** Differentially expressed gene (DEG) analysis was performed with DESeq2 (v1.44.0) in R (v4.4.0). Genes with $\log_2$ fold change absolute values ≥ 1 and adjusted p-value (padj) < 0.05 were considered differentially expressed with statistical significance. Volcano plots were generated using *ggplot2* version 4.0.0 comparing vehicle (High viral load) vs. uninfected (left) and favipiravir vs. uninfected (right) in the liver **(A)** and brain **(B)**.
