## Supplementary Tables for "Integrated *in vivo* and transcriptomic analyses of lethal Oropouche virus infection reveal suppression of pathogenic host responses by antiviral therapy"

| Gene name | Primers sequences (5' to 3') | Reference |
| --- | --- | --- |
| OROVS-F | TCCGGAGGCAGCATATGTG | [1] |
| OROVS-R | ACAACACCAGCATTGAGCACTT | |
| OROVS-Probe | (FAM)-CATTTGAAGCTAGATACGG-MGB(NFQ) | |
| Actb -F | GTGCTATGTTGCCCTGGACT | [2] |
| Actb -R | GCTCGTTGCCAATGGTGATG | |
| CXCL10 -F | GCCATTCATCCACAGTTGACA | [3] |
| CXCL10 -R | CATGGTGCTGACAGTGGAGTCT | |
| IL-6 - F | CCTGAAAGCACTTGAAGAATTCC | [3] |
| IL-6 - R | GGTATGCTAAGGCACAGCACACT | |
| IL-10 - F | GTTGCCAAACCTTATCAGAAATGA | [4] |
| IL-10 - R | TTCTGGCCCGTGGTTCTCT | |
| ISG15 - F | AAAGCCTACAGCCATGACCT | [5] |
| ISG15 - R | TTAGTCAGGGGCACCAGGAA | |
| RSAD2 -F | CGTGAGCATCGTGAGCAATG | [6] |
| RSAD2 -R | TGCACCACTTCCTCAGCTTT | |
| TNF-α - F | GGAGTGGCTGAGCCATCGT | [3] |
| TNF-α - R | AGCTGGTTGTCTTTGAGAGACATG | |

**S1 Table. List of primers used in this study.** MGB = minor-groove-binding; NFQ=non-fluorescent quencher

| Group | Sample ID | RNA Integrity number (RIN)/TapeStation | Reads | Yield(Gbases) | Q30 (%) |
|---|---|---|---|---|---|
| Mock (Liver) | LM1 | 8.6 | 19,539,342 | 5.8618 | 94.02 |
| | LM2 | 8.8 | 19,238,777 | 5.77163 | 93.76 |
| | LM3 | 8.8 | 19,153,207 | 5.74596 | 93.82 |
| Vehicle (Liver) | LV1 | 8.5 | 32,647,818 | 9.79435 | 94.61 |
| | LV2 | 8.7 | 44,821,940 | 13.44658 | 94.52 |
| | LV3 | 8.5 | 44,531,729 | 13.35952 | 94.56 |
| Treatment (Liver) | LT1 | 8.8 | 19,830,676 | 5.9492 | 94.03 |
| | LT2 | 7.8 | 18,192,060 | 5.45762 | 93.72 |
| | LT3 | 8.3 | 20,356,726 | 6.10702 | 93.70 |
| Mock (Liver) | BM1 | 8.8 | 35,577,954 | 10.67339 | 97.38 |
| | BM2 | 8.8 | 26,837,991 | 8.0514 | 93.13 |
| | BM3 | 8.3 | 18,362,293 | 5.50869 | 93.39 |
| Vehicle (Liver) | BV1 | 8.2 | 31,954,024 | 9.58621 | 94.05 |
| | BV2 | 8.2 | 22,236,817 | 6.67105 | 93.48 |
| | BV3 | 8.4 | 33,539,832 | 10.06195 | 94.28 |
| Treatment (Liver) | BT1 | 8.5 | 19,866,906 | 5.96007 | 93.78 |
| | BT2 | 8.7 | 20,131,326 | 6.0394 | 95.11 |
| | BT3 | 8.4 | 20,294,218 | 6.08827 | 93.58 |

**S2 Table. Summary of quality of isolated RNA and raw data statistics of the next generation sequencing related to Fig. 4.**
